# HIV-malaria co-infection and its determinants among patients attending antiretroviral treatment clinic in Zaria, North-Western Nigeria

**DOI:** 10.1101/588855

**Authors:** Shafi’u Dahiru Gumel, Abdulrasul Ibrahim, Adebola Tolulope Olayinka, Muhammed S. Ibrahim, Muhammad Shakir Balogun, Afara Dahiru, Ikeoluwapo Ajayi, Olufemi Ajumobi, Isiyaku Ahmadu, Abubakar Song, Asma’u Ibrahim Maifada, Habibu Abdullahi

**Affiliations:** Nigeria Field Epidemiology and Laboratory Training Program, Abuja, Nigeria; Department of Medical Microbiology, Ahmadu Bello Teaching Hospital, Zaria, Kaduna State, Nigeria; Nigeria Centre for Disease Control, Abuja, Nigeria; Department of Community Medicine, Faculty of Medicine, Ahmadu Bello University Zaria, Kaduna State, Nigeria; Department of Nursing, College of Nursing and Midwifery Hadejia, Jigawa State, Nigeria; Department of Epidemiology and Medical Statistics, Faculty of Public Health, College of Medicine, University of Ibadan, Oyo State, Nigeria; National Tuberculosis and Leprosy Treatment Centre, Saye, Kaduna State, Nigeria

**Keywords:** Determinants of HIV-malaria co-infection, malaria prevention, knowledge and practice of malaria prevention

## Abstract

**Introduction:** Malaria and HIV are two important global public health problems. Together, they cause more than two million deaths each year. In sub-Saharan Africa alone, more than 29 million people are living with HIV/AIDS and about 70% of population is at risk to malaria infection. Nigeria accounts for about a quarter of the global malaria cases and tenth of the global HIV cases. Recent theories suggested possibilities of high occurrence of HIV-malaria co-infection wherever there is geographical overlap of the two diseases. We therefore conducted this study to determine the prevalence of HIV-malaria co-infection and its determinants in a malaria endemic setting.

**Methods:** We conducted a cross-sectional study. Two hundred and sixty-two clients attending antiretroviral treatment (ART) clinic in Zaria, Kaduna State were enrolled between February and April 2018 using systematic sampling technique. Questionnaires were administered to collect information on respondents’ personal characteristics as well as their knowledge, perception and practices on malaria prevention. Venous blood samples were collected and analyzed for malaria parasite, viral load, CD4, and FBC using Giemsa stained light microscopy, COBAS TaqMan equipment, BD FACS™ flow cytometer, and Sysmex haematology analyser respectively. Descriptive and inferential statistics were performed, predictors of HIV-malaria co-infection were ascertained at multivariate analysis.

**Results:** Median age of the respondents was 33 years, 52% were females, 65% were married, 65% were employed, 57% lived in urban residence, and 34% had tertiary education. The prevalence of malaria co-infection among HIV patients was found to be 22.9%. Significant risk factors for the co-infection were high HIV viral load (aOR= 3.30, C.I = 1.15-9.45), being co-infected with TB (aOR= 5.60, C.I = 1.34-23.33), poor knowledge of malaria infection (aOR= 3.12, C.I = 1.27-7.72) and poor practice of malaria prevention (aOR= 13.30, C.I = 4.88-36.23).

**Conclusion:** The level of occurrence of malaria among HIV infected patients in this setting calls for attention. We recommended that health education on malaria should be a priority in malaria control programme; the programmes for control of HIV, malaria and TB should collaborate to ensure integrated service delivery and that people living with HIV/AIDS should be given special consideration for malaria prevention.

## Introduction

Human Immunodeficiency Virus (HIV) and malaria are major public health problems which are endemic in Nigeria (1,2). The HIV infection weakens the human immune system (3) making an individual more prone to other infections like malaria (4,5). Malaria on the other hand is caused by *Plasmodium*, a single cell microorganism that invades red blood cells, with greater propensity for severity and death in the presence of immunosuppression (5). It has also been reported that co-infection of HIV with malaria increases morbidity and mortality from HIV (6).

Risk of contracting malaria and developing severe disease varies between individuals and populations (7). These include children under 5 years of age (8), pregnant women (4), and immune-compromised patients such as those with HIV/AIDS (6). According to the World Health Organization (WHO), malaria and HIV combined, cause more than two million deaths each year (4). This is more so when the two diseases bring about common consequences such as anaemia (5,9–11).

There were an estimated 36.9 million people living with HIV (PLWHIV) in 2014 (1). Sub-Saharan Africa is the most affected region contributing about 66% (25.8 million) and Nigeria had about four million people living with HIV in 2014 (1).

In 2017, 219 million malaria cases and 435, 000 malaria related deaths were reported globally (12). Nigeria contributes about 23% of the global malaria cases and around 250,000 Nigerian children die from malaria every year (13). It is also estimated that 97% of Nigerian population is at risk of malarial infection (13).

There are reports that HIV and malaria co-infection may occur in any area that has high prevalence of the two infections (14,15) and that much possibilities of deleterious interaction between the two diseases exist in the co-infected patients (5,6,14). Some complications, such as anaemia, which are common to both HIV and malaria are also likely to be worse with the co-infection (16).This study was designed to determine the prevalence of malaria parasitaemia among HIV-positive clients in malaria endemic setting and to identify the determinants of this co-infection.

## Methods

### Study area

This study was conducted in Kaduna State, north-west Nigeria. In 2016, Kaduna state with HIV prevalence of 9.2% was ranked 3^rd^ among states with highest HIV burden in Nigeria (17). There are over 1,000 primary healthcare (PHC) facilities to cater for the health services of the residents. The HIV clinic at the National Tuberculosis and Leprosy Treatment Centre (NTBLTC) provides, alongside patient care, a comprehensive health education to the patients. This helps patient with knowledge about HIV and malaria infections, including prevention, treatment, and other aspect of care. It also equipped patients with tools that will enable them to participate more actively in decisions regarding their medical care. The area has two seasons, the rainy season (April–November) and the dry season (December–March). Malaria is endemic in the state but transmission is usually higher towards the end of the rainy season.

### Study design

A cross-sectional study was conducted among 262 consented persons living with HIV/AIDS (PLHIV) receiving treatment at the National Tuberculosis and Leprosy Training Center, Saye, Zaria, Kaduna State. The minimum sample size of 262 for the study was determined using 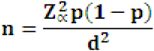 (18); taking prevalence (p) of HIV-malaria co-infection = 21% (19), confidence interval 95% and margin of error as 5%. Systematic sampling technique was used at a sampling interval of 16 (4,350/262) in a sampling frame of 4,350 hospital records. We included patients who had not received any anti-malaria drugs for one-month period, had tested positive for HIV, and who were neither severely ill nor mentally retarded.

### Data collection

Structured, interviewer administered questionnaire was used to collect data for this study. The questionnaire had five sections namely, Section A: Demographic information, Section B: Clinical information, Section C: Knowledge of malaria infection, Section D: Perception on malaria infection, and Section E: Practice of malaria prevention and control. The questionnaire was developed from earlier studies related to HIV and malaria; it was pre-tested before used and trained research assistants were used to administer the questionnaires.

The questionnaire had twelve questions on knowledge of malaria infection, six on perception on malaria infection and eight on practices of malaria prevention and control. Grading of these three parameters was done by assigning one mark for each correct answer and for multiple choices each correct response has one mark. A total of fifty, six, and sixteen marks were respectively allocated to knowledge, perception and practices. Fifty percent was considered as the cut-off point; in each category therefore, ≥ 50% marks was considered as good and < 50% marks as poor.

### Laboratory Examination

Two EDTA vacutainer tubes were used to collect 15 ml blood sample from each respondent; 10 ml and 5 ml separately for HIV viral load and the remaining laboratory tests respectively. From the 5 ml sample, thick and thin blood smears were made on grease-free slides and stained with Giemsa to determine species of malaria parasites and parasite density according to the earlier published protocol (20). Parasite densities were estimated by counting the number of malaria parasites against leukocytes using the following formula: 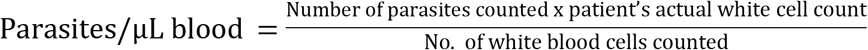 (21). All stained slides were examined by microscopy and read by two competent microscopists using 100 power fields under oil immersion. Polymerase Chain Reaction (PCR) using COBAS TaqMan equipment was employed to estimate the viral load of each respondent, and BD FACSPresto™ machine was used to estimate the CD4 count of each respondent adhering to the equipment manufacturer instructions.

### Statistical analysis

All data were entered into Microsoft excel, cleaned and imported into Epi Info version 7 statistical software for analysis. Univariate analysis including descriptive statistics like percentages and frequencies of the distribution of all variables of interest was performed. Chi-square test was conducted to determine the association between categorical variables i.e. association between HIV and malaria co-infection status in relation to age group, gender, place of residence, education level, employment status, ARV regimen, treatment duration, viral load level, CD_4_ count, TB status and knowledge, perception and practice of malaria infection prevention. Logistic regression model was conducted. Odds-ratios (OR) and adjusted odds-ratios (aOR) with 95% confidence interval (CI) were used to measure the strength of associations. All tests were two-tailed and P value < 0.05 was considered statistically significant.

### Ethical consideration

Ethical approval for the study was obtained from the Research and Ethics Committee of NTBLTC, Zaria. Written informed consent was obtained from the respondents.

## Results

The median age of the respondents was 32.50 years (Range: 3-74 years). One hundred and thirty-six respondents (51.9%) were females, 170 (64.9%) were married, 149 (56.9%) lived in urban residence, 89 (34.0%) had tertiary education, 115 (43.9%) were formally employed and 134 (51.2%) were of the Hausa-Fulani ethnic group (Table 1).

**Table 1:**
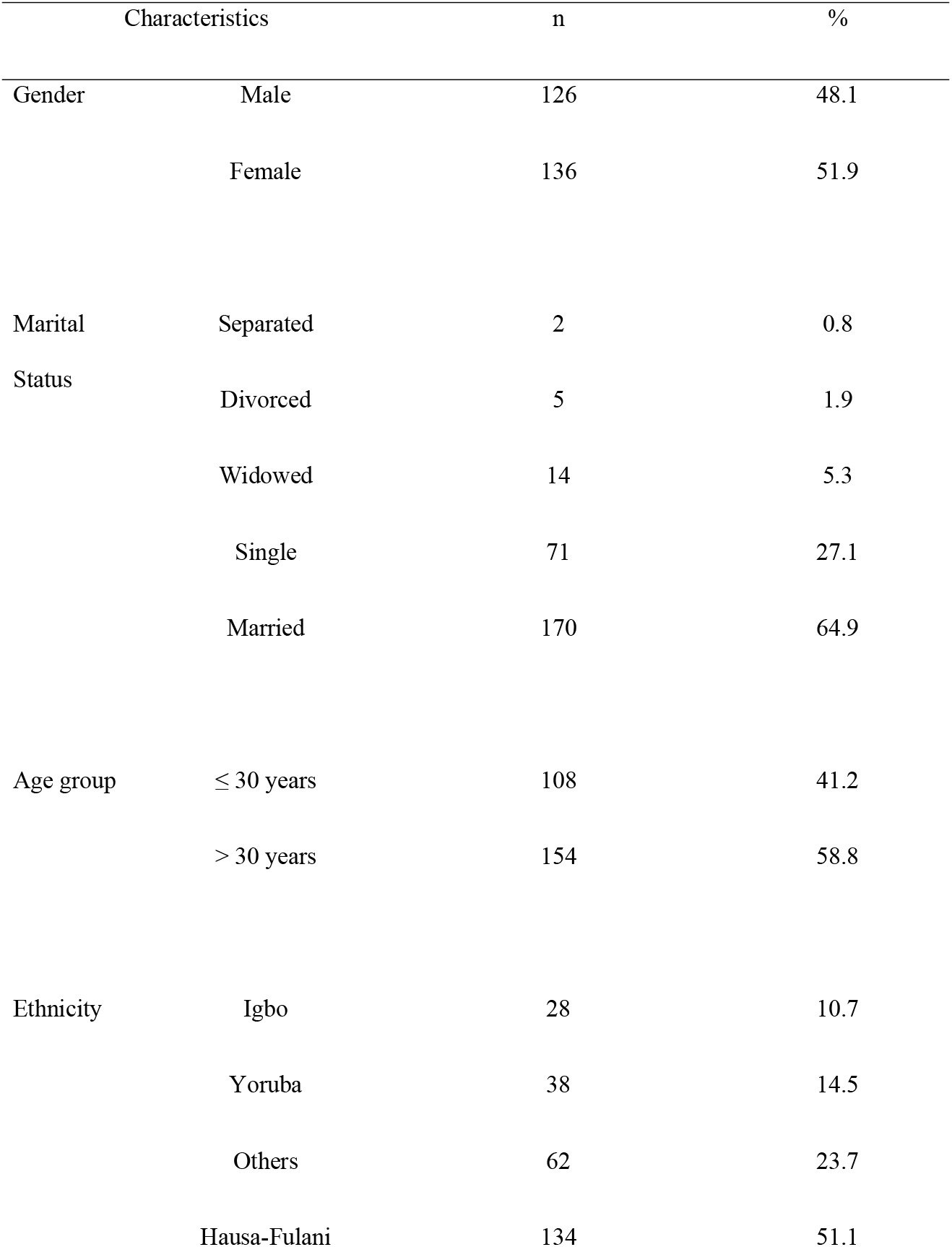
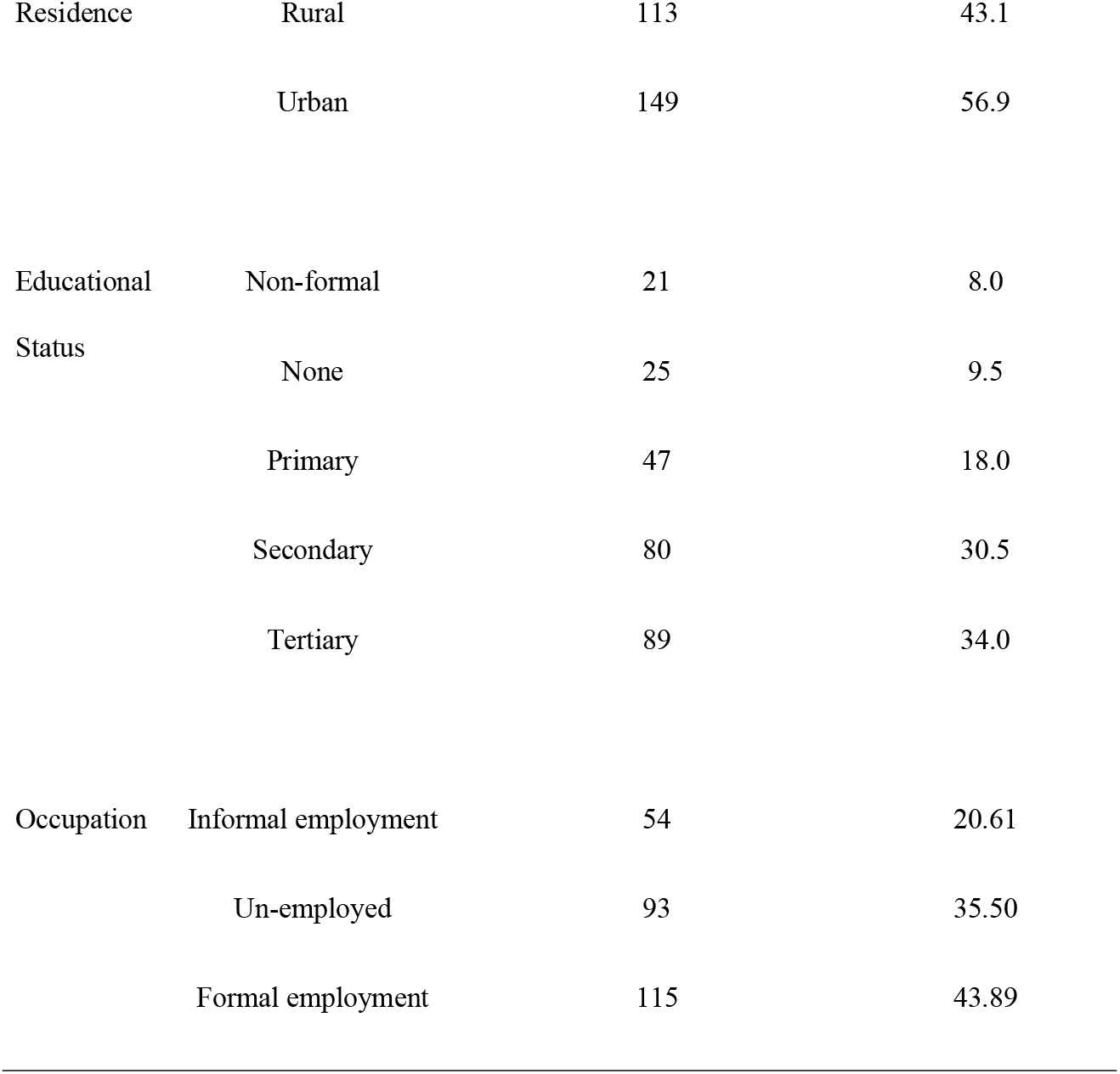
Socio-demographic characteristics of the respondents, NTBLTC Zaria, 2018 [N = 262]

| Characteristics |  | n | % |
| --- | --- | --- | --- |
| Gender | Male | 126 | 48.1 |
|  | Female | 136 | 51.9 |
| Marital Status | Separated | 2 | 0.8 |
|  | Divorced | 5 | 1.9 |
|  | Widowed | 14 | 5.3 |
|  | Single | 71 | 27.1 |
|  | Married | 170 | 64.9 |
| Age group | $\leq 30$ years | 108 | 41.2 |
| | $> 30$ years | 154 | 58.8 |
| Ethnicity | Igbo | 28 | 10.7 |
|  | Yoruba | 38 | 14.5 |
|  | Others | 62 | 23.7 |
|  | Hausa-Fulani | 134 | 51.1 |
| Residence | Rural | 113 | 43.1 |
|  | Urban | 149 | 56.9 |
| Educational<br>Status | Non-formal | 21 | 8.0 |
|  | None | 25 | 9.5 |
|  | Primary | 47 | 18.0 |
|  | Secondary | 80 | 30.5 |
|  | Tertiary | 89 | 34.0 |
| Occupation | Informal employment | 54 | 20.61 |
|  | Un-employed | 93 | 35.50 |
|  | Formal employment | 115 | 43.89 |

Two hundred and seventeen (82.8%) of the respondents had good knowledge about malaria infection and prevention, 172 (65.6%) had good perception and 145 (55.3%) had good practice. Of the total respondents, proportion of males with good knowledge, perception and practice was 107 (40.8%), 85 (32.4%), and 70 (26.7%) respectively (Fig 1).

**Figure 1:**
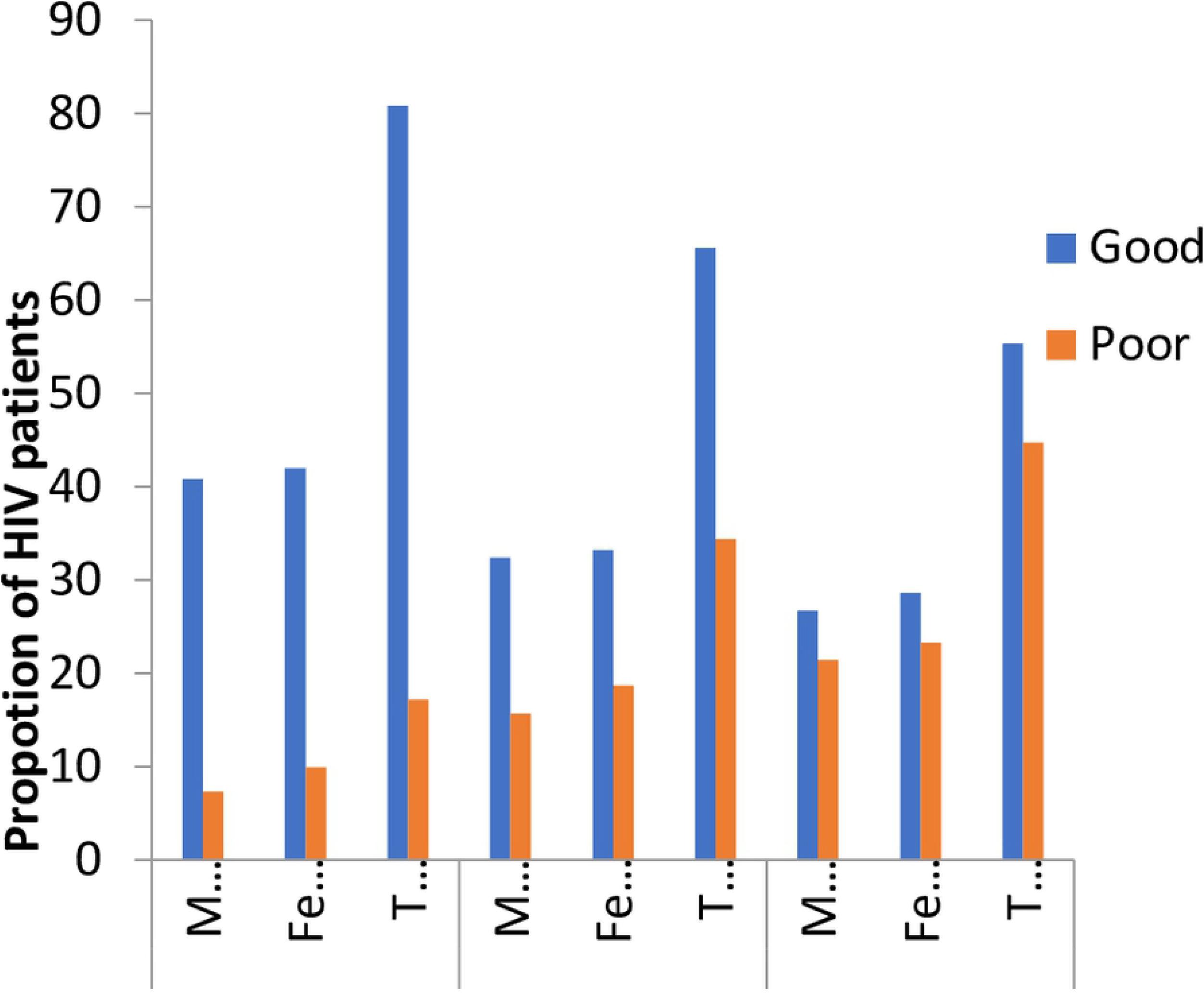
Knowledge, Perception, and Practice of the Respondents on Malaria Prevention by sex, NTBLTC Zaria, 2018

The prevalence of malaria co-infection among HIV patients was found to be 22.9%. On bivariate analysis, there was statistically significant association between malaria parasitaemia and residence, educational status, occupation, viral load, CD4 count, TB status and knowledge of malaria infection, and practice of malaria infection prevention and control (Table 2). However, on multivariate analysis only viral load, TB status, knowledge of malaria infection, and practice of malaria infection prevention and control were independently associated with malaria parasitaemia (Table 3).

**Table 2:** Risk Factors of HIV-Malaria co-infection, NTBLTC Zaria, 2018

| Characteristics |  | Malaria status |  | OR | 95% C.I |
| --- | --- | --- | --- | --- | --- |
|  |  | Positive | Negative |  |  |
| Gender: | Female | 31 | 95 | 1.20 | 0.67-2.14 |
|  | Male | 29 | 107 |  |  |
| Marital Status | Single | 41 | 129 | 1.22 | 0.66-2.26 |
|  | Married | 19 | 73 |  |  |
| Age group | > 30 years | 36 | 118 | 1.07 | 0.59-1.92 |
|  | ≤ 30 years | 24 | 84 |  |  |
| Residence | Rural | 34 | 79 | 2.04 | *1.14-3.65 |
|  | Urban | 26 | 123 |  |  |
| Formal Education | No | 23 | 23 | 4.84 | *2.46-9.53 |
|  | Yes | 37 | 179 |  |  |
| Formal employment | No | 48 | 99 | 4.16 | *2.09-8.30 |
|  | Yes | 12 | 103 |  |  |
| ARV Regimen | Second line | 14 | 37 | 1.36 | 0.68-2.72 |
|  | First line | 46 | 165 |  |  |
| Treatment Duration | > 5 years | 37 | 112 | 1.29 | 0.72-2.33 |
|  | ≤ 5 years | 23 | 90 |  |  |
| Viral load | High | 15 | 14 | 4.26 | *1.91-9.51 |
|  | Low | 42 | 167 |  |  |
| CD<sub>4</sub> Count | High | 32 | 75 | 1.94 | *1.08-3.46 |
|  | Low | 28 | 127 |  |  |
| TB Status | Positive | 10 | 10 | 3.84 | *1.51-9.73 |
|  | Negative | 50 | 192 |  |  |
| Knowledge | Poor | 24 | 21 | 5.75 | *2.90-11.41 |
|  | Good | 36 | 181 |  |  |
| Perception | Poor | 26 | 64 | 1.65 | 0.91-2.98 |
|  | Good | 34 | 138 |  |  |
| Practice | Poor | 52 | 65 | 13.70 | *6.15-30.51 |
|  | Good | 8 | 137 |  |  |

\*Variables significant at 5% level of significance

**Table 3:** Factors Independently Associated with HIV-Malaria co-infection, NTBLTC Zaria, 2018

| Characteristics | aOR | 95% C.I. |
| --- | --- | --- |
| Place of residence (Rural/Urban) | 1.38 | 0.59-3.22 |
| Formal Education (No/Yes) | 1.90 | 0.73-4.98 |
| Formal Employment (No/Yes) | 1.99 | 0.74-5.37 |
| Viral load (High/Low) | 3.30 | *1.15-9.45 |
| CD<sub>4</sub> Count (Low/High) | 1.92 | 0.87-4.24 |
| TB Status (Yes/No) | 5.60 | *1.34-23.33 |
| Knowledge (Poor/Good) | 3.12 | *1.27-7.72 |
| Perception (Poor/Good) | 2.23 | 0.93-5.33 |
| Practice (Poor/Good) | 13.30 | *4.88-36.23 |

\*Variables significant at 5% level of significance
aOR: Adjusted OR

## Discussion

In this study, the prevalence of malaria co-infection among individuals with HIV infection is 22.9%; this was slightly higher than 18.9% reported from Anambra, southeastern Nigeria(22) and 21% from Jos, northcentral Nigeria(19). The slight difference could be due to differing malaria burden between these 3 geo-political zones of the country. The proportion of HIV patients reported to be co-infected with malaria by the present and other studies indicates high occurrences of this phenomenon. In an attempt to explain the mechanism, it was indicated in 2014 that opsonizing antibodies and phagocytosis were significantly reduced in HIV-infected individuals(23); and these processes are very vital in body’s fight against malaria.

In this study, the prevalence of the co-infection does not differ significantly between males and females and is not associated with age. This agrees with findings from a study in Kano, North-West Nigeria(24) but contrary to another one from Jos North-Central Nigeria(19). This could be explained by the fact that the study conducted in Jos was among subjects that were mainly urban residents unlike that of the present study which comprises both urban and rural dwellers. In urban areas, there seems to be some differences in knowledge and practice of malaria infection prevention between the different gender and age groups unlike in rural areas where everything seems to be the same, this may explain the disparity in findings of these studies.

Risk factors for HIV-malaria co-infection were also studied in this study; viral load, TB status, knowledge of malaria infection, and practice of malaria infection prevention were found to be associated with malaria co-infection in HIV patients. Other factors that may confound or modify the association were place of residence, educational status, and occupation.

These findings agree with several similar studies and theories. For instance, even though people residing in areas of stable malaria transmission develop humoral and cell-mediated immunity to malaria parasite; it was shown that this immunity can be altered in HIV-infected persons and could influence the frequency and course of malaria infection(25). This could therefore explain the association of the viral load with malaria infection where by the risk increases with increase in viral load.

With regards to TB, it is well documented that HIV increases the risk of TB infection and vice versa (26,27); the current understanding of the human immune response to malaria, HIV, and TB make one to expect that TB infections could influence the clinical outcome of malaria in HIV clients. Infection with TB also weakens the body immune system increasing vulnerability to malaria infection.

This study also showed that rural dwellers had the highest proportion of those with HIV-malaria co-infection. This could be attributed to lack or inadequate materials for malaria prevention, such as insecticide treated nets, in the rural areas compared to urban areas (28), leading to much exposure of rural dwellers to consistent bite by mosquitoes and therefore malaria. This is supported by the earlier report of Bassiouny and Al-Maktari in 2005 that rural areas have environmental conditions more favorable to transmission of the disease than urban areas (29).

In our study also, poor knowledge of malaria infection and poor practice of malaria prevention were significant risk factors to malaria co-infection in HIV patients. In a systematic review of knowledge, attitudes and beliefs about malaria among the South Asian population, Krishna *et al*, shows that various measures to prevent malaria may exist but the success of these measures depends on the knowledge of, access to and utilization of services, as well as a combination of users’ behaviors and healthcare access and quality issues (30). This could help in explaining the concurrent findings of high knowledge of the subjects and high prevalence of malaria co-infection among the respondents in this study because it is not all knowledge that will translate to good practice.

## Conclusion

From findings of this study, the prevalence of malaria co-infection among HIV patients in Zaria, Kaduna State is of public health concern. Having high HIV viral load, being co-infected with TB, having poor knowledge of malaria infection and poor practice of malaria infection prevention are significant risk factors for HIV-malaria co-infection. While place of residence, education level and occupation may modify or have a confounding effect on the association of these risk factors with the co-infection rate.

We therefore recommend that, because of their vulnerability to malaria, people living with HIV/AIDS should be considered among priority group for any malaria intervention; government at federal, state and local government levels should consider improving education of people as a means of tackling the co-infection; programmes for control of the two diseases and that of TB should collaborate to ensure integrated service delivery; these programmes should consider rural dwellers among priority groups; and additional research on interactions between the two diseases should be prioritised.

## Acknowledgment

The authors wish to appreciate the staff of National TB Reference laboratory Saye-Zaria, who provided the much-needed support and a conducive atmosphere for the successful conduct of this study.

